# Making the best of a bad job? Chick mortality and flexible female brood care in Snowy Plovers

**DOI:** 10.1101/2019.12.19.880856

**Authors:** Krisztina Kupán, Tamás Székely, Medardo Cruz-López, Keeley Seymour, Clemens Küpper

## Abstract

Offspring desertion represents a trade-off between current and future reproductive success. Its timing is crucial as the termination of parental care has profound consequences for the fitness of the parents and their offspring. However, the decision process involved with termination of care is still poorly understood. Snowy Plovers *Charadrius nivosus* show highly flexible brood care with females either deserting the brood early or providing care for an extended period. Deserting females often quickly remate and start a new breeding attempt. Using a dynamic modelling framework, we investigated the decision-making process for continuation or termination of care by females over a seven-year period. The length of female care increased over the season likely reflecting lower re-mating opportunities for deserting females late in the season. Present brood size, assessed daily during the brood care period, was strongly related to the length of female care: females were more likely to stay and care for larger than for smaller broods. Chick death and desertion frequently coincided, suggesting that poor offspring condition served as a trigger for female desertion. Overall, deserting females had a similar number of fledglings to caring females. This suggests that for many females, desertion was not a strategy to escape the shackles of monogamy and secure higher reproductive success through sequential polygamy. Rather, most deserting females made the best of a bad job when conditions were poor and their continued presence did not make a difference for the survival of their young. We conclude that when making the decision to continue or terminate care, Snowy Plover females monitor the condition of their offspring closely and adjust their care flexibly to the value and needs of their young.

## Introduction

Parental care increases parental fitness by improving offspring growth, condition and ultimately survival (Klug & Bonsall, 2014). At the same time parental care adds major costs to reproduction that may compromise adult survival and/or future fecundity of the parent (Buzatto, Requena, Martins, & Machado, 2007; Clutton-Brock, 1991; Gross & Sargent, 1985; Smith & Wootton, 1995; Zink, 2003). In particular, multiple breeders of long-lived species face a trade-off between improving the prospects of the current brood through care and the reduction of their own future reproductive success and/or survival through extended care (Ackerman, Eadie, Yarris, Loughman, & McLandress, 2003; Magrath & Komdeur, 2003; Trivers, 1972; James N Webb, Houston, McNamara, & Székely, 1999; Westneat & Sargent, 1996; Williams, 1966).

One possible way of escaping the strains of parental duty is premature care termination through offspring desertion. Whilst originally considered as maladaptive or ‘abnormal behaviour’ (Fujioka, 1989; Hrdy, 1999), theoretical and experimental studies have shown that offspring desertion is often beneficial for the parents as it can increase lifetime reproductive success (Clutton-Brock, 1991; Maynard Smith, 1977; Smith & Wootton, 1995; Székely, Webb, Houston, & McNamara, 1996; Trivers, 1972; Ward, Cotter, & Kilner, 2009; James N Webb et al., 1999). In order to make an adaptive decision over desertion, parents should consider five key parameters: 1) the needs of the current offspring, 2) the value and prospects of the current brood, 3) re-mating opportunities, 4) their own condition, and 5) behaviour of the other parent in biparental systems (Houston, Székely, & McNamara, 2005; Kelly & Kennedy, 1993; Székely, 1996; J. N. Webb, Székely, Houston, & McNamara, 2002).

First, the needs of the current brood may depend on the environment, in which the offspring will grow up. In a harsh environment one parent may not be able to raise the young alone whereas in a benign environment the offspring may fare as well with one parent as with both, leaving one parent free to desert (Amat, Fraga, & Arroyo, 1999a; Beissinger & Snyder, 1987; Blanken & Nol, 1998; Eldegard & Sonerud, 2009; Kosztolányi et al., 2009; Magrath & Komdeur, 2003; Székely et al., 1996).

Second, the value and prospects of the current brood is primarily determined by brood size and offspring condition. Brood size is the most obvious predictor of desertion with parents usually caring longer for larger than for smaller broods (e.g. (Beissinger & Snyder, 1987; Fujioka, 1989; Steinhart, Dunlop, Ridgway, & Marschall, 2008; Ward et al., 2009)). Again, brood prospects may chiefly depend on environmental conditions, thus parents need to assess offspring survival with or without its contribution (Steinhart et al., 2008).

Third, the deserting partner has to adjudge its re-mating opportunities. Re-mating opportunities determine who gains more from deserting the family and who gains more from care (Balshine-Earn & Earn, 1998; Eldegard & Sonerud, 2009; Fujioka, 1989; Keenleyside, 1985; Pilastro, Biddau, Marin, & Mingozzi, 2001; Roulin, 2002; Thomson et al., 2014). Recent studies have shown that reproductive behaviour is often tied to sex ratios of the adult population (Grant & Grant, 2019; Kokko & Jennions, 2008; Liker, Freckleton, & Székely, 2013) with biased sex ratios altering potential for sequential polygamy in each sex. In that case, the rarer sex has a higher opportunity for polygamy and is more likely to desert the brood (Eberhart-Phillips et al., 2018).

Fourth, the parent should consider its future reproductive prospects. Parents need to assess their own condition and survival prospect in this process. When parental care is depleting energy reserves or increasing the risk of predation, desertion may improve the prospect of adult survival and enable them to reproduce again (Currie et al., 2001; Jamieson, 2012; Osorno, 1999).

Fifth, a deserting parent will leave dependent offspring behind. The deserter needs to consider the ability and behaviour of the remaining parent, who will be left with the young alone. When the remaining parent is able to fully compensate for the lack of the second carer continued care by both parents is not necessary (Fujioka, 1989; Harrison, Barta, Cuthill, & Székely, 2009; Osorno & Székely, 2004; Roulin, 2002; Székely et al., 1996; Ward et al., 2009). One parent may be enough when care is non-depreciable (i.e. parental effort does not increase substantially with the number of offspring, (Clutton-Brock, 1991; Smith & Wootton, 1995), or the young reached an age where they require little parental care. In these cases, desertion is more likely to happen although sexual conflict between the parents over care may be high (Beissinger, 1987).

Importantly, all of these factors are changing over the brood care period. Furthermore, they are often intertwined and not independent from each other. It is essential to understand the relationship between these factors in order to establish the motive behind and the adaptivity of early termination of care. For example, the brood size might decrease over the brood care period apparently decreasing the value of the current brood. However, the value for the parents often also depends on the re-mating opportunities: if re-mating opportunities are high parents may be better off prioritizing future reproduction over their current offspring. Contrarily, if re-mating opportunities are low, it may be more advantageous for the parent to continue caring for their current young. Often the timing of desertion may hint towards the underlying cause of desertion. Desertion early in the season points towards prioritizing higher reproductive success by taking advantage of high re-mating opportunities whereas desertion late in the season may point towards prioritizing their own survival (Currie et al., 2001; Osorno, 1999). When desertion happens during an early stage of care, the young are still vulnerable and the remaining parent might not be able to fully compensate. In this case, early termination of care will only be adaptive if the gains from additional matings are higher than the losses incurred in the deserted brood (Eldegard & Sonerud, 2009; Ezaki, 1988; van Dijk, 2009).

It is, however, often not clear whether low survival prospects of the offspring are cause or consequence of desertion. In the former case, the deserting parent may essentially abandon a sinking ship as continued care will not improve the offspring survival prospects. Instead, the deserter tries to make the best of a bad situation by starting a fresh mating attempt to offset its initial low reproductive success. In the latter case, the deserter already takes into account lower future survival of the deserted offspring but expects to obtain an overall higher reproductive success through re-mating than when remaining with the first brood.

Plovers (Charadriinae) are small shorebirds with high variation and flexibility in parental care (e.g. (Eberhart-Phillips et al., 2018; Vincze et al., 2017). They typically lay small clutches of with two to four eggs imply that reproductive success can only increase through producing multiple clutches (or double brooding) (Blomqvist, Wallander, & Andersson, 2001). Precociality of the offspring has facilitated the evolution of flexible brood care systems that may feature bi- or uniparental brood care (Eberhart-Phillips, 2019; Houston et al., 2005; Székely & Reynolds, 1995; Thomas & Székely, 2005). The frequency of desertion varies not only between species but also between and within populations indicating extraordinary flexibility (Eberhart-Phillips et al., 2018). Several predictors of the length of biparental care have been identified previously including initial brood size (also called as “current brood size” (Székely & Cuthill, 2000; Ward et al., 2009)), biotic environment (such as population density and/or predation pressure (Amat et al., 1999a; Kosztolányi, Székely, Cuthill, Yilmaz, & Berberoğlu, 2006)) and abiotic environment (e.g. temperature, (AlRashidi et al., 2010)). Desertion is strongly linked to re-mating opportunities that are influenced by adult sex ratios: typically, the rarer sex has higher re-mating opportunities than the more common sex and therefore deserts primarily (Amat et al., 1999a; Carmona-Isunza et al., 2017; Eberhart-Phillips et al., 2018; Eberhart-Phillips et al., 2017; Parra, Beltrán, Zefania, Dos Remedios, & Székely, 2014; Stenzel et al., 2011; Székely, Cuthill, & Kis, 1999). Moreover, the length of biparental care increases over the season as re-mating opportunities diminish (Amat et al., 1999a; Kosztolányi et al., 2009; Kosztolányi et al., 2006; Székely & Cuthill, 1999; Székely et al., 1999).

In many plover populations, desertion appears positively related to chick mortality (Amat, Fraga, & Arroyo, 1999b; Cruz-López et al., 2017; Székely, 1996; Székely & Cuthill, 1999; Székely et al., 1996; Székely & Williams, 1995). We previously reported that the majority of deserted Snowy Plover *Charadrius nivosus* families were characterised by low chick survival, which might imply that brood desertion is maladaptive (Cruz-López et al., 2017). However, whether desertion is cause or consequence of reduced chick survival remained unclear. Following up on these results, we investigated the phenology and fitness consequences of flexible brood care in Snowy Plovers breeding at Bahía de Ceuta, Mexico (aka "Ceuta”). Using the fates of 268 families collected over seven breeding seasons we assessed the adaptive value and the dynamics of female brood care and offspring desertion. We used a recently developed dynamic modelling method (Schmidt, Walker, Lindberg, Johnson, & Stephens, 2010) to identify both static and dynamic predictors of female desertion. We considered present brood size (the brood size on a specific day) and brood age as dynamic variables that represent changing values and needs of the brood. Specifically, we 1) describe the pattern of female brood care strategies in this population, 2) test which social and environmental variables predict the length of female care and, 3) examine the temporal association of chick mortality and termination of female care. With this we explore whether and how chick mortality influences the decision of the females to continue or terminate care.

## Methods

### Field work and data collection

We studied breeding Snowy Plovers at Ceuta, a coastal wetland in Northwest Mexico between April and July from 2006 to 2012. About 50–100 Snowy Plover pairs nested on tidal salt flats of an abandoned salt extraction site surrounded by mangrove forests and agricultural fields annually during the study period (Cruz-López et al., 2017; Eberhart-Phillips et al., 2017; Plaschke, Bulla, Cruz-Lopez, del Ángel, & Küpper, 2019). We monitored Snowy Plover families daily using the methodology described in ref. (Székely, Kosztolanyi, & Küpper, 2008). For further details see Supplementary Material.

### Data processing and statistical analysis

We collected brood attendance data for a total of 367 broods. From these, 12 (3.3%) did not have hatching date information, 37 (10%) had two or less observations, in 7 (2%) broods the male deserted or disappeared and in a further 43 (11.7%) broods the female deserted during an unobserved period larger than three days. After removing these broods, we included 268 broods for further analyses. Against a strong population decline throughout the study period, brood numbers varied annually (Figure S1) (Cruz-López et al., 2017).

All broods in the final dataset were followed from the hatching date of the first chick (day 0) until fledging (day 25) or complete failure. We assumed that all parents were present with the clutch until hatching as incubation in Snowy Plovers is biparental (Vincze et al., 2017; Vincze et al., 2013). We calculated the duration of female and male care for each brood in days since hatching of the first chick. As we performed brood manipulations such as cross-fostering of freshly hatched chicks and eggs during some years that led to altered brood sizes, we used for each brood the hatching date of the chicks the parents cared for.

For each bird, we defined the last day of presence as the mid-point between the last observation day with the family and the first day it was absent permanently. When the gap between observations of uni- and biparental care was an odd number of days, we used the ceiling value as the length of biparental care. We considered temporarily absence, i.e. birds that were subsequently re-sighted with the family as ‘present’. The maximum duration of care was 26 days (from day 0 until day 25). With this method we calculated the length of presence for family members, which represents survival of hatchlings and duration of care for either parent. When repeating all analyses with a reduced data set that excluded broods with estimated female desertion dates we obtained qualitatively similar results (results available on request).

### I. Female brood care

In this study we focused on female care as this is the most flexible in this Snowy Plover population whereas males almost always invariably care (Cruz-López et al., 2017). We considered four scenarios to describe the termination of female care. 1) *Desertion:* the female left the brood whilst at least one chick was still attended by the male. 2) *Brood failure:* either both parents were observed without the chicks, or, when the male was then seen alone within three days of the last observation of the brood attended by both parents. 3) *Full term care:* the female stayed until at least one chick fledged. 4) *Unfinished observations:* unfledged broods with unknown fate that were attended by the female at the last observation.

To assess the fitness consequences of female brood care strategies, we compared the seasonal reproductive success and breeding effort of females that for the first clutch either provided full term care or deserted and then re-mated locally (Supplementary Material).

### II. Predictors of the length of female brood care

As a dynamic measure for the length of female care we calculated a novel response variable: *probability of care*. *Probability of care* is analogous to *daily nest survival*, an established unbiased measure in ecology to understand fundamental mechanisms of population dynamics (Converse, Royle, Adler, Urbanek, & Barzen, 2013; Schmidt et al., 2010). *Probability of care* refers to the probability that a female will care on a certain day given the probability of care on the previous day. We adopt this measure to assess the drivers for termination of parental care as it allows us to assess not only static but also dynamic predictors that are changing over the care period. We analysed the effects of seven biologically meaningful predictor variables on the *probability of care*: *hatching date*, *male tarsus length*, *male condition, female condition, chick hatching condition* were static predictors and *brood age* and *present brood size* were dynamic predictors (see detailed description of the different predictors in the Supplementary Material). *Present brood size*, the brood size on a certain day, may provide a more precise estimate on the effect of brood size than initial brood size on care. Note that *present brood size* differs from the commonly used ‘current brood size’ (Székely & Cuthill, 2000; Ward et al., 2009), which refers to the initial brood size of the current breeding attempt. By contrast, in our study *present brood size* may change during the care period, i.e. it decreases with chick mortality. Basing our models on present rather than initial brood sizes allows us to investigate the female decision to continue or terminate care without assuming the ‘Concorde fallacy’ (Dawkins & Carlisle, 1976), i.e. that past investment instead of future prospect has the strongest impact on the parental care decision (Ackerman et al., 2003).

We set up a binomial generalized linear mixed model to estimate the effect of the predictors on *probability of care* using a Markov-chain Monte Carlo (MCMC) algorithm in a Bayesian framework (Korner-Nievergelt et al., 2015). We used STAN (Stan Development Team, 2018) through the R packages *rstan*, *rstanarm* (Stan Development Team, 2018) and *arm* (Gelman & Hill, 2007). We fitted two random intercepts: *year* and *female ID* since females can have multiple breeding events within and between years. We modelled *probability of care* of each female with Bernoulli errors.

The likelihood to continue care for a given day was formulated as

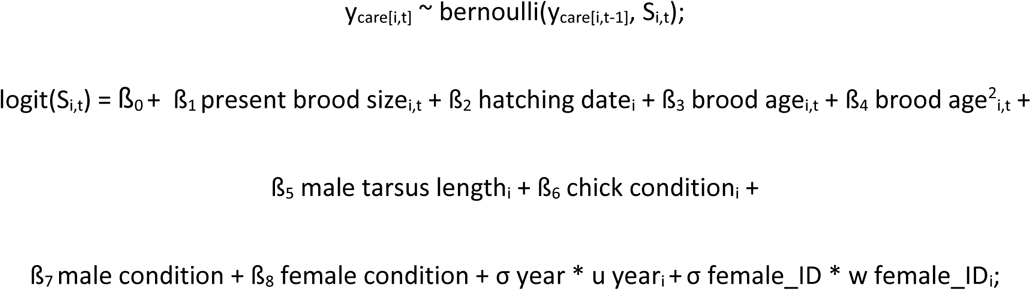

where *y*_*care*_ represents female care as binary (‘1’ for care and ‘0’ for no care) with dimensions *i* as the number of broods (n = 268) and *t* as the length of the brood care period (0 to 25 days). *S* is the daily care continuation probability, which relates to brood and day specific predictors of the model. All fixed variables were z-transformed with mean = 0 and standard deviation (sd) = 1. We used normal priors with mean = 0 and sd = 5 on fixed effects, normal priors (mean = 0, sd = 1) on random effects and Cauchy (0,5) priors on the sigma parameters (year, female_ID).

Since STAN currently does not accept missing values for the estimations, we substituted missing values with the population average for *male tarsus length* and *condition* variables. Importantly, the model can distinguish between different ‘end of care’ events. The last value the model considered for a certain brood was always the last observation day, i.e. the first day the female did not provide care anymore in deserted and failed broods, therefore had the value of 0. Alternatively, for broods with unknown fate or full term biparental care the value was 1 and referred to the last day the female still provided care (Korner-Nievergelt et al., 2015). We obtained samples of the posterior distribution of the model parameters from five independent Markov chains with 4000 iterations each. We discarded the first 2000 values of the burn-in period of each chain and then calculated posterior parameter estimates from the remaining iterations. For analytics and visual inspection to determine convergence of the model we used *shinystan* package (Stan Development Team, 2018). All models had fully converged with the 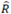 values = 1 (Gelman, Carlin, Stern, & Rubin, 2004). For visualization of the results we used the *bayesplot* (Gabry & Mahr, 2017) and *MCMCvis* packages in R (Youngflesh, 2018).

### III. Termination of care and chick mortality

In this analysis we examined the relationship between female termination of care and chick mortality. First, we tested whether females were more likely to *terminate care* on days when one or all chick died than when the brood size did not change. We included all broods belonging to the categories desertion, brood failure and full term care excluding unfinished broods. Second, we examined whether females were more likely to *desert* on days when chick mortality happened. For this, we only considered deserted broods and tested whether desertion occurred randomly or coincided with chick loss. We included all broods for which the last pre- and first post-desertion observations were no more than three days apart. Deteriorating chick condition, because of starvation, injuries or diseases, may precede chick loss by one or two days. Females may be able to anticipate chick death and leave such broods already before the actual reduction of brood size as we often observed chicks moving sluggishly and lagging behind the family shortly before they disappeared (CK & MCL, personal observations).

We used co-occurrence models (conditional logistic regression) to test whether female desertion and chick mortality co-occurred more frequently than expected by chance. We coded desertion (i.e. the change in probability of care) as ‘1’ and chick mortality as ‘1’, and continued female care and no chick loss as ‘0’. We added *Nest ID* as a grouping variable for the model. We used the ‘clogit’ function of the *survival* R package for the analysis (Therneau & Lumley, 2015). For all statistical analyses we used R version 3.5.3. (R Development Core Team, 2010).

## Results

### I. Female brood care

Among the 268 Snowy Plover broods examined, desertion was the most frequent terminal event of female care with 197 (74%) of all broods deserted. Desertion typically happened during early brood care, i.e. in 93% of the deserted broods the female left within ten days after hatching (Figure S2). The peak of desertion occurred two days after hatching (Figure S2). In stark contrast, only a minority of six (2%) females cared for the brood until fledging. Thirteen (5%) broods failed before the female deserted. Finally, for 52 (19%) broods the terminal fate was not known and these females were still seen with the brood at the last observation.

Comparing reproductive success and breeding effort revealed that caring and deserting females had similar reproductive success but deserting females required considerably higher breeding effort than caring females to achieve this (Supplementary Material, Figure S4).

### II. Predictors of the length of female brood care

*Hatching date*, *present brood size* and *brood age^2^* were all associated with female *probability of care* (Figure 1a). Early in the season females terminated care quickly to desert the brood (Figure 1b). By the end of the season females continued to care longer and *probability of care* increased from 0.44 (95% CrI: 0.21-0.66) to 0.96 (95% CrI: 0.92-0.98) over the season. The p*robability of care* for females caring for large broods were higher than those for females caring for small broods (Figure 1c). Mean *probability of care* for the broods with one chick was 0.64 (95% CrI: 0.42–0.8) and increased by approximately 0.04 for every additional chick to 0.81 (95% CrI: 0.65–0.9) for a five chick brood, the largest brood size in our sample. *Brood age^2^* also showed a clear association with length of female care. Rates of *probability of care* decreased from 0.81 (95% CrI: 0.74-0.87) at hatching to 0.72 (95% CrI: 0.56-0.82) at a brood age of 9 days and then increased to 0.93 (95% CrI: 0.7-0.99) for females that stayed until the age of 25 days (Figure 1d).

**Figure 1.**
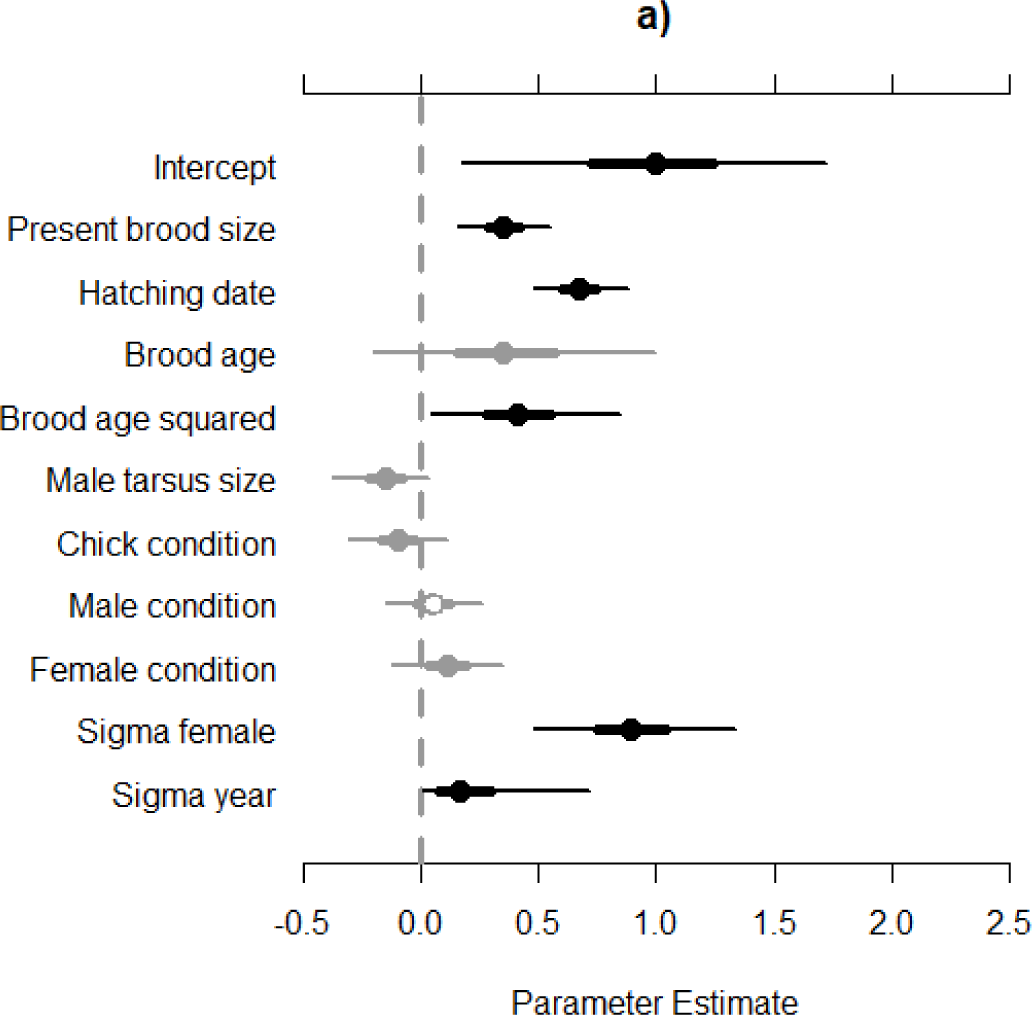
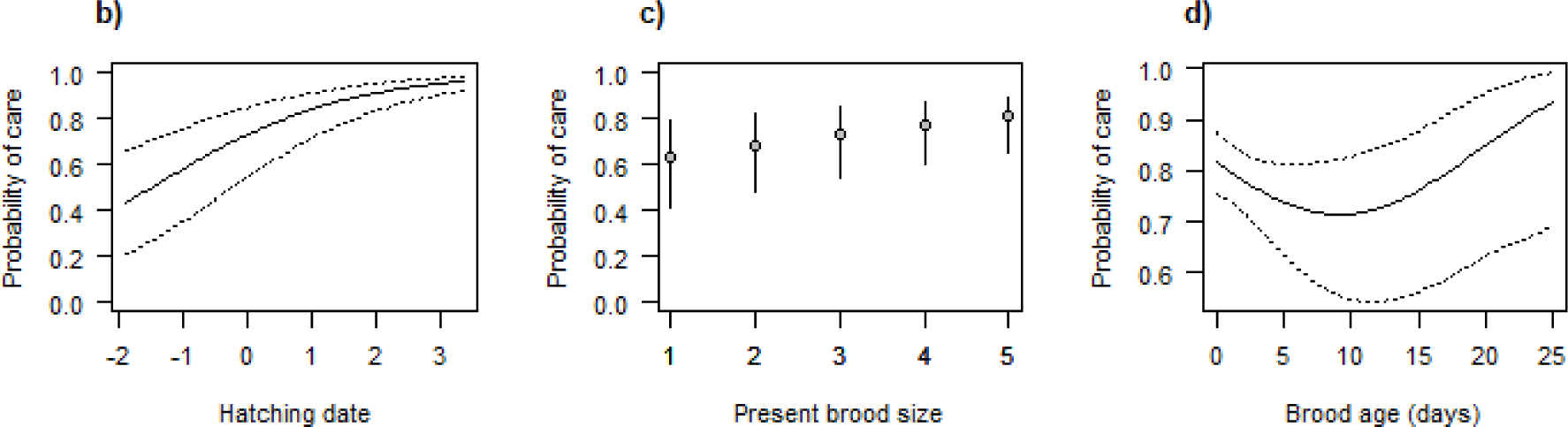
Predictors of the *probability of female care* in 268 Snowy Plover broods at Ceuta when all other traits are kept at the mean. (a) Summary of all model predictors. *Sigma_female* and *Sigma_year* are random effects. Dots represent means, thick lines standard deviations and thin lines are the 95% credibility intervals (CrI). Black symbols indicate no CrI overlap with zero, for grey symbols CrIs overlap with zero. (b,c,d) Details for fixed effects with a statistically clear impact on the length of female care: *hatching date* (standardized to the annual mean) (b), *present brood size* (c), and *brood age* (d). Continuous lines refer to mean estimates and dotted lines represent the credibility intervals (b, d), circles represent the mean and continuous lines represent CrIs (c).

Neither *male tarsus size* nor any of the *condition* variables were statistically clearly related to the *probability of care* as CrIs overlapped with zero (Figure 1a). Similarly, brood manipulations had no clear effect on female *probability of care* (Supplementary Material, Figure S5a). The analysis without the manipulated broods did not change the results qualitatively (Supplementary Material, Figure S5b).

### III. Termination of care and chick mortality

In 48 (22%) broods the female terminated care on the very day a chick died (Figures 2 & 3). The conditional logistic regression model showed a clear association between chick mortality and female termination of care (p < 0.001). The co-occurrence between termination of care and chick mortality was stronger for broods with two or more chicks as nearly all females tending one-chick broods deserted before the chick had died or fledged. Furthermore, in all of the broods with three or more chicks the female deserted once a chick died (Table S1, Figure S6). Restricting the analysis to include only those broods that were deserted (i.e. excluding one chick broods where desertion and mortality cannot co-occur by definition) resulted in an even stronger co-occurrence between female care termination and chick mortality (24%, p < 0.001). Chick survival was significantly lower in broods, in which co-occurrence happened than in those broods in which the two events did not co-occur (p < 0.001, Supplementary Material, Figure S7).

**Figure 2.**
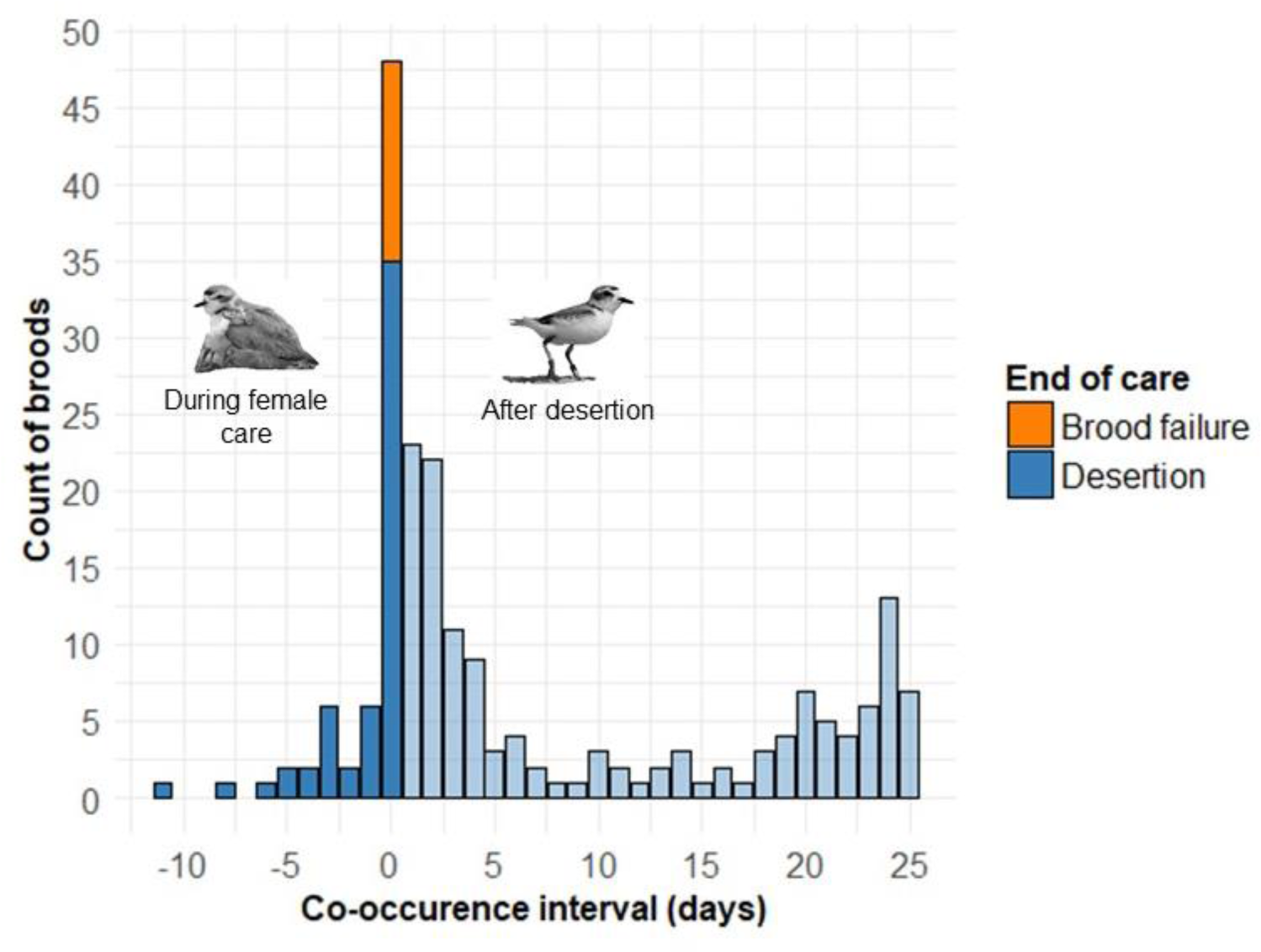
Co-occurrence of chick mortality and female care termination in 182 Snowy Plover broods with known dates for chick death or fledging. The diagram shows the interval between the most recent chick death and female termination of care within broods. Mortality and care termination coincide on the same day when the co-occurrence interval is zero. Negative values refer to mortality whilst the female still cared (dark blue). Positive values refer to cases when chick mortality happened after the female had deserted (light blue).

## Discussion

How do parents decide whether to continue care or desert their offspring? Our analysis of female brood care in Snowy Plovers suggests that females constantly assess the needs, value and survival prospects of their offspring whilst they provide care. This ability is a precondition for an adaptive decision about whether or not to continue care. Deserting and re-mating Snowy Plovers attempt to maximize their reproductive success by rapid divorce after hatching as divorced females produce a higher number of chicks than females that retain their mates (Halimubieke et al., 2019). Yet, we confirmed that desertion does not necessarily translate into producing more fledglings (Cruz-López et al., 2017). Our findings suggest that full term care providing females reach a similar reproductive success (measured as number of fledglings) as deserting and locally re-mating females but in significantly shorter time. Moreover, occasionally, caring females will still have enough time left to establish a new clutch and may hence increase their reproductive success further. The low number of locally re-mating females suggests that many deserting females disperse further to find new partners. Some of these dispersing females may reach higher reproductive success than locally re-mating females (Halimubieke et al., 2019). However, the breeding effort of dispersers is higher than that of caring or locally re-mating females as breeding dispersal will take up further time.

When analysing variables associated with the length of female care, we found both static and dynamic predictors that related to the probability that females will continue care. The dynamic modelling approach allowed us to analyse the consequences of changes in these predictors on individual decisions of females. Our study identified three key predictors for the length of female care. First, the hatching date of a brood was positively associated with the length of female brood care. In early broods, females cared for a shorter time and deserted the broods faster than in late broods. This is consistent with desertion for sequential polygamy and has also been observed in many other plover populations (Amat et al., 1999a; Kosztolányi et al., 2009; Székely & Cuthill, 2000; Székely & Williams, 1994; Warriner, Warriner, Page, & Stenzel, 1986). Re-mating opportunities are decreasing with every day, so fast desertion will maximize the reproductive potential of early breeding females whereas late breeding females may be better off providing care. Deserting females in Ceuta likely benefit from high re-mating opportunities due to a male-biased adult sex ratio that translates into a male-biased operational sex ratio (Carmona-Isunza et al., 2017; Eberhart-Phillips, 2017). In contrast to females, males have low remating opportunities and are therefore probably better off remaining with the brood and providing care for the offspring.

Second, we found that the *present brood size* had a strong effect on the probability of female care. Females were less likely to stay when brood size decreased during the brood care period. This suggests that for females the value of the brood is not fixed and determined by initial brood size and/or the breeding season (Székely & Cuthill, 2000) but rather is assessed regularly over the brood care period. Therefore, Snowy Plover females avoid the ‘Concorde fallacy’ (Ackerman et al., 2003; Armstrong & Robertson, 1988; Dawkins & Carlisle, 1976). That survival prospects of the brood, particularly chick mortality, eminently affected the female’s decision whether to continue or terminate care is shown by the close temporal association between chick mortality and desertion. The co-occurrence of chick mortality and desertion was much stronger than expected by chance. Furthermore, broods with co-occurrence were characterised by lower chick survival than broods without co-occurrence. This suggest that females are able to assess the survival prospects of the brood. Chick death seemed to be an important trigger for females to terminate brood care albeit our data did not allow us to firmly determine whether the mortality necessarily preceded desertion. Instead some females may already desert the brood if one chick is in bad condition due to sickness or starvation and, therefore, likely to die regardless of female care. We found that for broods with two or more chicks the co-occurrence of female care termination and chick mortality was particularly high. At the same time nearly all broods with one chick were deserted by the female before the chick had died or fledged. This suggests that one-chick broods do not have enough reproductive value for the female to stay and care.

Third, we found that the probability of female care changed with brood age. Snowy Plover chicks are most vulnerable during the first three days of their lives (Colwell, Hurley, Hall, & Dinsmore, 2007) and the peak of desertion fell into this period. We observed an initial decline in the probability to care until the age of nine days. This is consistent with our expectations based on the needs of the brood. As chicks grow older, they improve their thermoregulation and require less brooding, one of the most time consuming care behaviours (Székely and Cuthill, 2000). Hence, the presence of the second carer becomes less important. Interestingly, after the age of nine days the relationship changed and the probability to continue to care increased until fledging. In precocial species such as the Snowy Plover, chick survival typically increases with age (Colwell et al., 2007; Cruz-López et al., 2017). For females that already cared for an extended period, care continuation may also be the best option as the value of the current brood is high, especially at the end of the season when the re-mating opportunities are low (Figure 1c, (Székely & Cuthill, 2000)).

Similar to other studies, condition and size related predictors were not clearly associated with the length of female care (Amat et al., 1999b; Amat, Visser, Perez-Hurtado, & Arroyo, 2000; Székely & Williams, 1995). As desertion rates declined with season, desertion does not seem to be a response to energy depletion as in other species (Currie et al., 2001; Gratto‐Trevor, 1991; Osorno, 1999).

Parental brood care in plovers mainly consists of thermoregulation, warning chicks of predators and defending them from attacks of competitors (Carmona-Isunza et al., 2017). However, chick survival is also strongly dependent on the availability of high quality habitat with invertebrate food which declines over the season (Cruz-López et al., 2017; dos Remedios, Lee, Burke, Székely, & Küpper, 2015; Kosztolányi et al., 2006). In a stochastic environment flexible care behaviour that follows the changing value and needs of their brood can be highly advantageous. In Snail Kites *Rostrhamus sociabilis*, for example, one parent tends to desert the brood once a single parent can care for the chicks alone. In this system chick survival is generally low, therefore, parents do not compromise offspring survival by desertion. Rather desertion happens once the needs of the brood reached the ‘monoparental threshold’ either due to brood size reduction or to advanced brood age (Beissinger, 1986, 1990). In other studies the value of broods has been shown to be important for desertion. Female ducks tended to desert their clutches once the number of predated eggs reached a certain proportion (Ackerman et al., 2003). Our results suggest that for many Snowy Plovers females in this population desertion may have two different motives: i) increase the reproductive success through sequential polygamy or ii) make the best of a bad job. Both scenarios are represented by distinct clusters in Figure 3. Early breeding females may utilize the good environmental conditions and desert to increase their reproductive success by producing a second clutch quickly without jeopardising the survival of the deserted brood (Osorno, 1999). Similar to the ‘monoparental threshold’ (Beissinger, 1990), the value of the current brood is high whilst the needs for care are low. By contrast, to make the best of a bad job, females desert when their parental effort cannot prevent chick mortality. This is the main reason for desertion in our population. Chick starvation and flooding, which are the main threats for offspring failure in the Ceuta population (Cruz-López et al., 2017; Plaschke et al., 2019), cannot be mitigated by the parents. After the death of one or more chicks the value of the brood will become too low for the female to continue care (similar to the ‘value threshold’, (Ackerman et al., 2003)). Although the needs of the brood are still high, they cannot be fulfilled by the female. Therefore, having an additional mating attempt will allow females to compensate for the reproductive losses. Consequently, extended biparental care is only expected when both the needs and the value of the brood are high and parental care fulfils the offspring needs, e.g. through protection.

**Figure 3.**
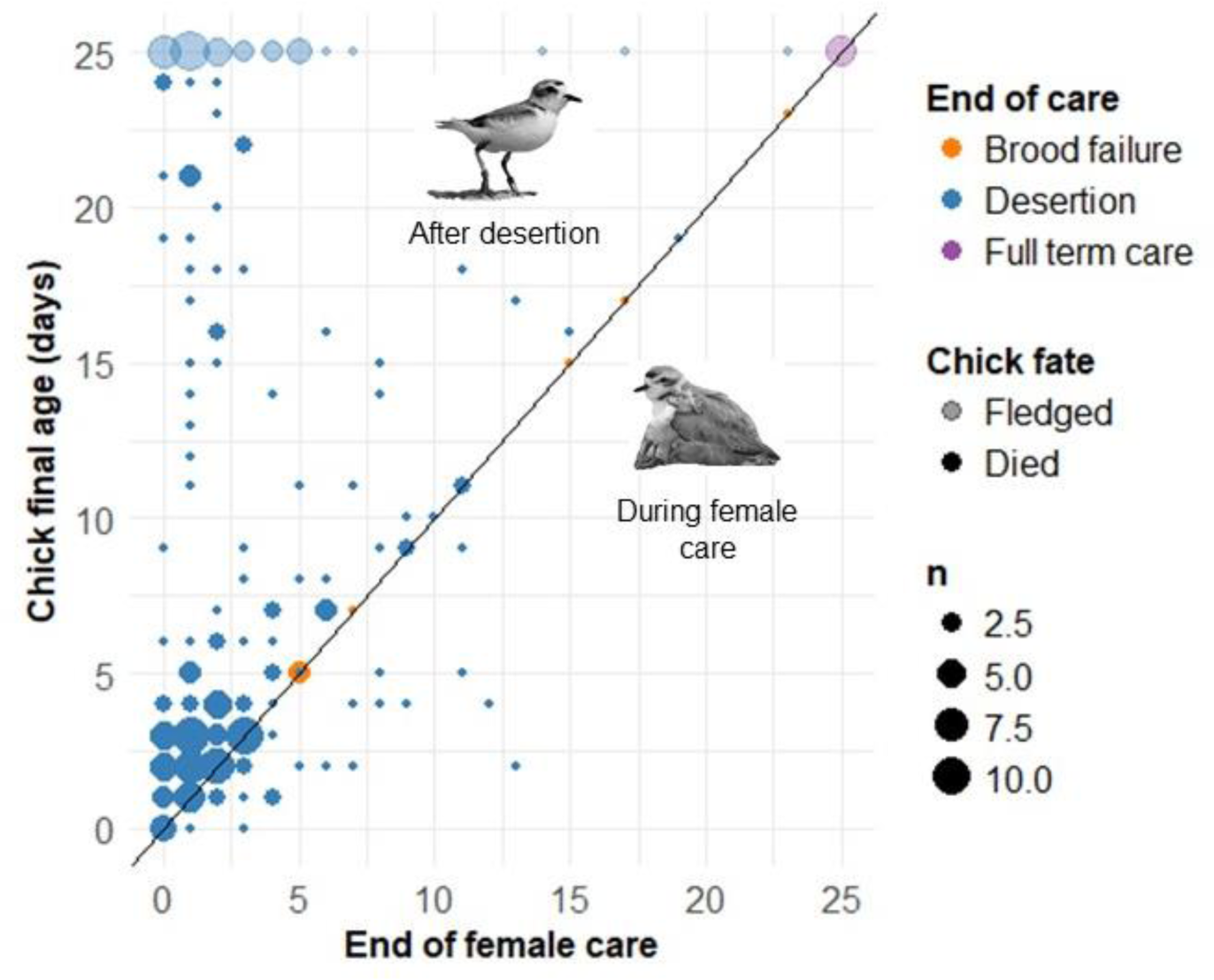
Chick survival in relation to length of female care for 182 Snowy Plover broods with known dates for chick death or fledging. Only one chick per brood, whose fate was most closely associated with the end of female care, is plotted. Dot sizes refer to number of broods. Coordinate (0,0) indicates the day of hatching and hence the start of female brood care. The diagonal line indicates co-occurrence of chick fate (died or fledged) and female care termination on the same day. Below the diagonal are broods in which the female continued to care after one chick had died, above the diagonal are broods where the female deserted before the first chick had died.

In conclusion, the examination of high resolution data of parental care and chick survival with a dynamic modelling approach revealed multiple facets of female desertion within a single species. Our analysis of female care in Snowy Plover shows a strong association between the probability of care and both, the present value of the current brood and future mating opportunities. The strong association of chick mortality and female care termination suggests that high re-mating opportunities provide females with a plan B to rescue some reproductive success when environmental conditions are poor. However, deserting females represent a heterogeneous group. They include some successful females that pursue polyandrous matings to multiply their reproductive success although most deserting females use sequential polyandry to compensate for low chick survival in their current brood. Importantly, our study demonstrates that the decision over care or desertion is dynamically changing and responds to both, the current needs and the value of the offspring.

## Acknowledgements

We thank Lydia Lozano-Angulo, Raul Quintero-Felix, Oscar Sánchez-Velázquez, Karla Alvarado-Castro, Wendy Rojas-Abreu and René Beamonte-Barrientos for help with fieldwork. Fränzi Korner-Nievergelt and Mihai Valcu provided valuable advice on the statistical analyses. A list of funding sources for fieldwork is provided on www.chorlito.org. K.K. and C.K. are supported by the Max Planck Society. T.S. was funded by a Royal Society Wolfson Merit Award (WM170050) and by the National Research, Development and Innovation Office of Hungary (ÉLVONAL KKP-126949, K-116310). The authors declare no conflict of interest.

## Authors’ contributions

T.S., K.S. and K.K. conceived the project. C.K. and M.C.L. provided the data. K.K. carried out the statistical analyses. K.K and C.K. wrote the manuscript with input from all authors.

## Data availability statement

All data and R codes will be made available on GitHub repository: https://github.com/kreastie/Plover-Desertion-Project

## References

Ackerman, J. T., Eadie, J. M., Yarris, G. S., Loughman, D. L., & McLandress, M. R. (2003). Cues for investment: nest desertion in response to partial clutch depredation in dabbling ducks. Animal Behaviour, 66(5), 871–883.

AlRashidi, M., Kosztolányi, A., Küpper, C., Cuthill, I. C., Javed, S., & Székely, T. (2010). The influence of a hot environment on parental cooperation of a ground-nesting shorebird, the Kentish plover *Charadrius alexandrinus*. Frontiers in Zoology, 7(1), 1. doi:10.1186/1742-9994-7-1

Amat, J. A., Fraga, R. M., & Arroyo, G. M. (1999a). Brood desertion and polygamous breeding in the Kentish Plover *Charadrius alexandrinus*. Ibis, 141(4), 596–607.

Amat, J. A., Fraga, R. M., & Arroyo, G. M. (1999b). Replacement clutches by Kentish Plovers. Condor, 101(4), 746–751.

Amat, J. A., Visser, G. H., Perez-Hurtado, A., & Arroyo, G. M. (2000). Brood desertion by female shorebirds: a test of the differential parental capacity hypothesis on Kentish plovers. Proceedings of the Royal Society of London Series B, 267(1458), 2171–2176.

Armstrong, T., & Robertson, R. J. (1988). Parental investment based on clutch value: nest desertion in response to partial clutch loss in dabbling ducks. Animal Behaviour, 36(3), 941–943.

Balshine-Earn, S., & Earn, D. J. (1998). On the evolutionary pathway of parental care in mouth–brooding cichlid fishes. Proceedings of the Royal Society of London. Series B: Biological Sciences, 265(1411), 2217–2222.

Beissinger, S. R. (1986). Demography, environmental uncertainty, and the evolution of mate desertion in the snail kite. Ecology, 67(6), 1445–1459.

Beissinger, S. R. (1987). Mate desertion and reproductive effort in the snail kite. Animal Behaviour, 35(5), 1504–1519.

Beissinger, S. R. (1990). Experimental brood manipulations and the monoparental threshold in snail kites. The American Naturalist, 136(1), 20–38.

Beissinger, S. R., & Snyder, N. F. R. (1987). Mate desertion in the snail kite. Animal Behaviour, 35(2), 477–487.

Blanken, M. S., & Nol, E. (1998). Factors affecting parental behavior in Semipalmated Plovers. The Auk, 166–174.

Blomqvist, D., Wallander, J., & Andersson, M. (2001). Successive clutches and parental roles in waders: the importance of timing in multiple clutch systems. Biological Journal of the Linnean Society, 74(4), 549–555. doi:10.1006/bijl.2001.0593

Buzatto, B. A., Requena, G. S., Martins, E. G., & Machado, G. (2007). Effects of maternal care on the lifetime reproductive success of females in a neotropical harvestman. Journal of Animal Ecology, 76(5), 937–945.

Carmona-Isunza, M. C., Ancona, S., Székely, T., Ramallo-González, A. P., Cruz-López, M., Serrano-Meneses, M. A., & Küpper, C. (2017). Adult sex ratio and operational sex ratio exhibit different temporal dynamics in the wild. Behavioral Ecology, 28(2), 523–532. doi:10.1093/beheco/arw183

Clutton-Brock, T. H. (1991). The evolution of parental care: Princeton University Press.

Colwell, M. A., Hurley, S. J., Hall, J. N., & Dinsmore, S. J. (2007). Age-related survival and behavior of Snowy Plover chicks. The Condor, 109(3), 638–647.

Converse, S. J., Royle, J. A., Adler, P. H., Urbanek, R. P., & Barzen, J. A. (2013). A hierarchical nest survival model integrating incomplete temporally varying covariates. Ecology and Evolution, 3(13), 4439–4447.

Cruz-López, M., Eberhart-Phillips, L. J., Fernández, G., Beamonte-Barrientos, R., Székely, T., Serrano-Meneses, M. A., & Küpper, C. (2017). The plight of a plover: Viability of an important snowy plover population with flexible brood care in Mexico. Biological Conservation, 209, 440–448.

Currie, D., Valkama, J., Berg, Å., Boschert, M., Norrdahl, K., Häninnen, M., … Hemminki, O. (2001). Sex roles, parental effort and offspring desertion in the monogamous Eurasian Curlew Numenius arquata. Ibis, 143(3), 642–650. doi:10.1111/j.1474-919X.2001.tb04892.x

Dawkins, R., & Carlisle, T. R. (1976). Parental investment, mate desertion and a fallacy. Nature, 262(5564), 131.

dos Remedios, N., Lee, P. L., Burke, T., Székely, T., & Küpper, C. (2015). North or south? Phylogenetic and biogeographic origins of a globally distributed avian clade. Molecular Phylogenetics and Evolution, 89, 151–159.

Eberhart-Phillips, L. J. (2017). Dancing in the moonlight: evidence that Killdeer foraging behaviour varies with the lunar cycle. Journal of Ornithology, 158(1), 253–262. doi:10.1007/s10336-016-1389-4

Eberhart-Phillips, L. J. (2019). Plover Breeding Systems. In The Population Ecology and Conservation of Charadrius Plovers (Vol. 53, pp. 65).

Eberhart-Phillips, L. J., Küpper, C., Carmona-Isunza, M. C., Vincze, O., Zefania, S., Cruz-López, M., … Krüger, O. (2018). Demographic causes of adult sex ratio variation and their consequences for parental cooperation. Nature Communications, 9(1), 1651. doi:10.1038/s41467-018-03833-5

Eberhart-Phillips, L. J., Küpper, C., Miller, T. E. X., Cruz-López, M., Maher, K. H., dos Remedios, N., … Székely, T. (2017). Sex-specific early survival drives adult sex ratio bias in snowy plovers and impacts mating system and population growth. Proceedings of the National Academy of Sciences, 114(27), E5474. doi:10.1073/pnas.1620043114

Eldegard, K., & Sonerud, G. A. (2009). Female offspring desertion and male-only care increase with natural and experimental increase in food abundance. Proceedings of the Royal Society B: Biological Sciences, 276(1662), 1713–1721.

Ezaki, Y. (1988). Mate desertion by male Great Reed Warblers Acrocephalus arundinaceus at the end of the breeding season. Ibis, 130(3), 427–437. doi:10.1111/j.1474-919X.1988.tb08817.x

Fujioka, M. (1989). Mate and nestling desertion in colonial little egrets. The Auk, 106(2), 292–302.

Gabry, J., & Mahr, T. (2017). bayesplot: Plotting for Bayesian models. R package version, 1.

Gelman, A., Carlin, J. B., Stern, H. S., & Rubin, D. (2004). Bayesian data analysis 2nd edn Chapman & Hall.

Gelman, A., & Hill, J. (2007). Data analysis using regression and multilevel/hierarchical models. New York: Cambridge University Press.

Grant, P. R., & Grant, B. R. (2019). Adult sex ratio influences mate choice in Darwin’s finches. Proceedings of the National Academy of Sciences, 116(25), 12373–12382. doi:10.1073/pnas.1903838116

Gratto-Trevor, C. (1991). Parental care in semipalmated sandpipers Calidris pusilla: brood desertion by females. Ibis, 133(4), 394–399.

Gross, M. R., & Sargent, R. C. (1985). The Evolution of Male and Female Parental Care in Fishes. American Zoologist, 25, 807–822.

Halimubieke, N., Valdebenito, J. O., Harding, P., Cruz‐López, M., Serrano-Meneses, M. A., James, R., … Székely, T. (2019). Mate fidelity in a polygamous shorebird, the snowy plover (Charadrius nivosus). Ecology and Evolution.

Harrison, F., Barta, Z., Cuthill, I., & Székely, T. (2009). How is sexual conflict over parental care resolved? A meta-analysis. Journal of Evolutionary Biology, 22(9), 1800–1812. doi:10.1111/j.1420-9101.2009.01792.x

Houston, A. I., Székely, T., & McNamara, J. M. (2005). Conflict between parents over care. Trends in Ecology & Evolution, 20(1), 33–38.

Hrdy, S. B. (1999). Mother nature: A history of mothers, infants, and natural selection.

Jamieson, S. E. (2012). Body mass dynamics during incubation and duration of parental care in Pacific Dunlins Calidris alpina pacifica: a test of the differential parental capacity hypothesis. Ibis, 154(4), 838–845. doi:10.1111/j.1474-919X.2012.01255.x

Keenleyside, M. H. (1985). Bigamy and mate choice in the biparental cichlid fish Cichlasoma nigrofasciatum. Behavioral Ecology and Sociobiology, 17(3), 285–290.

Kelly, E. J., & Kennedy, P. L. (1993). A dynamic state variable model of mate desertion in Cooper’s hawks. Ecology, 74(2), 351–366.

Klug, H., & Bonsall, M. B. (2014). What are the benefits of parental care? The importance of parental effects on developmental rate. Ecology and Evolution, 4(12), 2330–2351. doi:10.1002/ece3.1083

Kokko, H., & Jennions, M. D. (2008). Parental investment, sexual selection and sex ratios. Journal of Evolutionary Biology, 21, 919–948.

Korner-Nievergelt, F., Roth, T., Von Felten, S., Guélat, J., Almasi, B., & Korner-Nievergelt, P. (2015). Bayesian data analysis in ecology using linear models with R, BUGS, and Stan: Academic Press.

Kosztolányi, A., Javed, S., Küpper, C., Cuthill, I. C., Al Shamsi, A., & Székely, T. (2009). Breeding ecology of Kentish Plover *Charadrius alexandrinus* in an extremely hot environment. Bird Study, 56, 244–252.

Kosztolányi, A., Székely, T., Cuthill, I. C., Yilmaz, K. T., & Berberoğlu, S. (2006). Ecological constraints on breeding system evolution: the influence of habitat on brood desertion in Kentish plover. Journal of Animal Ecology, 75(1), 257–265. doi:10.1111/j.1365-2656.2006.01049.x

Liker, A., Freckleton, R. P., & Székely, T. (2013). The evolution of sex roles in birds is related to adult sex ratio. Nature Communications, 4, 1587–1587. doi:10.1038/ncomms2600

Magrath, M. J., & Komdeur, J. (2003). Is male care compromised by additional mating opportunity?. Trends in Ecology & Evolution, 18(8), 424–430.

Maynard Smith, J. (1977). Parental investment: A prospective analysis. Animal Behaviour.

Osorno, J. L. (1999). Offspring Desertion in the Magnificent Frigatebird: Are Males Facing a Trade-Off between Current and Future Reproduction?. Journal of Avian Biology, 30(4), 335–341. doi:10.2307/3677005

Osorno, J. L., & Székely, T. (2004). Sexual conflict and parental care in magnificent frigatebirds: full compensation by deserted females. Animal Behaviour, 68, 337–342.

Parra, J. E., Beltrán, M., Zefania, S., Dos Remedios, N., & Székely, T. (2014). Experimental assessment of mating opportunities in three shorebird species. Animal Behaviour, 90, 83–90.

Pilastro, A., Biddau, L., Marin, G., & Mingozzi, T. (2001). Female brood desertion increases with number of available mates in the rock sparrow. Journal of Avian Biology, 32(1), 68–72.

Plaschke, S., Bulla, M., Cruz-Lopez, M., del Ángel, G., & Küpper, C. (2019). Nest initiation and flooding in response to season and semi-lunar spring tides in a ground-nesting shorebird. Frontiers in Zoology, 16(1), 11.

R Development Core Team. (2010). R: A language and environment for statistical computing.. Retrieved from http://www.R-project.org

Roulin, A. (2002). Offspring desertion by double-brooded female barn owls (Tyto alba). The Auk, 119(2), 515–519.

Schmidt, J. H., Walker, J. A., Lindberg, M. S., Johnson, D. S., & Stephens, S. E. (2010). A general Bayesian hierarchical model for estimating survival of nests and young. The Auk, 127(2), 379–386.

Smith, C., & Wootton, R. J. (1995). The costs of parental care in teleost fishes. Reviews in Fish Biology and Fisheries, 5(1), 7–22.

Stan Development Team. (2018). RStan: the R interface to Stan.

Steinhart, G. B., Dunlop, E. S., Ridgway, M. S., & Marschall, E. A. (2008). Should I stay or should I go? Optimal parental care decisions of a nest-guarding fish. Evolutionary Ecology Research, 10(3), 351–371.

Stenzel, L. E., Page, G. W., Warriner, J. C., Warriner, J. S., Neuman, K. K., George, D. E., … Bidstrup, F. C. (2011). Male-skewed adult sex ratio, survival, mating opportunity and annual productivity in the Snowy Plover *Charadrius alexandrinus*. Ibis, 153, 312–322.

Székely, T. (1996). Brood desertion in Kentish Plover Charadrius alexandrinus: An experimental test of parental quality and remating opportunities. Ibis, 138(4), 749–755.

Székely, T., & Cuthill, I. C. (1999). Brood desertion in Kentish plover: the value of parental care. Behavioral Ecology, 10(2), 191–197.

Székely, T., & Cuthill, I. C. (2000). Trade-off between mating opportunities and parental care: brood desertion by female Kentish plovers. Proceedings of the Royal Society of London Series B, 267(1457), 2087–2092.

Székely, T., Cuthill, I. C., & Kis, J. (1999). Brood desertion in Kentish plover: sex differences in remating opportunities. Behavioral Ecology, 10(2), 185–190.

Székely, T., Kosztolanyi, A., & Küpper, C. (2008). Practical guide for investigating breeding ecology of Kentish plover Charadrius alexandrinus. Unpublished Report. University of Bath.

Székely, T., & Reynolds, J. D. (1995). Evolutionary transitions in parental care in shorebirds. Proceedings of the Royal Society of London Series B, 262(1363), 57–64.

Székely, T., Webb, J. N., Houston, A. I., & McNamara, J. M. (1996). An evolutionary approach to offspring desertion in birds. Current ornithology, 271–330.

Székely, T., & Williams, T. D. (1994). Factors affecting timing of brood desertion by female Kentish plovers *Charadrius alexandrinus*. Behaviour, 130, 17–28.

Székely, T., & Williams, T. D. (1995). Costs and benefits of brood desertion in female Kentish plovers *Charadrius alexandrinus*. Behavioral Ecology and Sociobiology, 37(3), 155–161.

Therneau, T. M., & Lumley, T. (2015). Package ‘survival’. R Top Doc, 128.

Thomas, G. H., & Székely, T. (2005). Evolutionary pathways in shorebird breeding systems: Sexual conflict, parental care, and chick development. Evolution, 59, 2222–2230.

Thomson, R. L., Pakanen, V.-M., Tracy, D. M., Kvist, L., Lank, D. B., Rönkä, A., & Koivula, K. (2014). Providing parental care entails variable mating opportunity costs for male Temminck’s stints. Behavioral Ecology and Sociobiology, 68(8), 1261–1272.

Trivers, R. L. (1972). Parental investment and sexual selection. In B. Campbell (Ed.),. Sexual selection and the descent of man 1871-1971 (pp. 136–179). Chicago: Aldine.

van Dijk, R. E. (2009). Sexual conflict over parental care in penduline tits. (Doctor of Philosophy). University of Bath, Bath, UK.

Vincze, O., Kosztolányi, A., Barta, Z., Küpper, C., Alrashidi, M., Amat, J. A., … Székely, T. (2017). Parental cooperation in a changing climate: fluctuating environments predict shifts in care division. Global Ecology and Biogeography, 26(3), 347–358. doi: doi:10.1111/geb.12540

Vincze, O., Székely, T., Küpper, C., AlRashidi, M., Amat, J. A., Ticó, A. A., … Kosztolányi, A. (2013). Local environment but not genetic differentiation influences biparental care in ten plover populations. PLOS One, 8(4), e60998. doi:10.1371/journal.pone.0060998

Ward, R. J., Cotter, S. C., & Kilner, R. M. (2009). Current brood size and residual reproductive value predict offspring desertion in the burying beetle Nicrophorus vespilloides. Behavioral Ecology, 20(6), 1274–1281.

Warriner, J. S., Warriner, J. C., Page, G. W., & Stenzel, L. E. (1986). Mating system and reproductive success of a small population of polygamous snowy plovers. Wilson Bulletin, 98(1), 15–37.

Webb, J. N., Houston, A. I., McNamara, J. M., & Székely, T. (1999). Multiple patterns of parental care. Animal Behaviour, 58(5), 983–993.

Webb, J. N., Székely, T., Houston, A. I., & McNamara, J. M. (2002). A theoretical analysis of the energetic costs and consequences of parental care decisions. Philosophical Transactions of the Royal Society of London. Series B: Biological Sciences, 357(1419), 331–340.

Westneat, D. F., & Sargent, R. C. (1996). Sex and parenting: the effects of sexual conflict and parentage on parental strategies. Trends in Ecology & Evolution, 11(2), 87–91.

Williams, G. C. (1966). Natural selection, the costs of reproduction, and a refinement of Lack’s principle. The American Naturalist, 100(916), 687–690.

Youngflesh, C. (2018). MCMCvis: Tools to Visualize, Manipulate, and Summarize MCMC Output. J. Open Source Software, 3(24), 640.

Zink, A. G. (2003). Quantifying the costs and benefits of parental care in female treehoppers. Behavioral Ecology, 14(5), 687–693.

