## Supplementary Material for "Making the best of a bad job? Chick mortality and flexible female brood care in Snowy Plovers"

**Field data collection**

We searched for nests and families by scanning the salt flats for incubating plovers with binoculars and scopes from a mobile hide or a car. When a nest was found, we recorded the location with a hand-held GPS device (Garmin, USA) and established laying and hatching dates based on the floating technique assuming a 25 days incubation period (Plaschke, Bulla, Cruz-Lopez, del Ángel, & Küpper, 2019). We caught parents of the nests whilst on incubation duty with a funnel trap and marked them with a unique colour-metal-ring combination consisting of three colour rings and a numbered metal ring. We revisited the nests approximately every two to four days until we heard the chicks calling inside the eggs and afterwards daily to capture, measure and mark chicks before they leave the nest scrape (Cruz-López et al., 2017; Dos Remedios, Székely, Küpper, Lee, & Kosztolányi, 2015). We marked the chicks with a metal and a single colour ring. This allowed us to follow the individual fates of the chicks. We captured most broods at hatching at or in the vicinity of the nest scrape. A small number of chicks (<5%) from unknown nests were caught together with the attending parent at opportunistic encounters in the field. For these cases estimated the hatching date based on tarsus length of the chicks (Dos Remedios et al., 2015). We took body measurements and a blood sample of adults and chicks for molecular sex identification. We re-sighted families if possible at least once every two days until the brood had reached an age of five days and approximately every three days until the brood age reached 25 days at which brood age Snowy Plovers can be independent (Cruz-López et al., 2017). At each sighting we observed broods until each previously attending parent had been seen or for 15 minutes to identify all family members present (Székely & Cuthill, 1999). If a bird was not seen during this period we recorded it as 'missing' for that day. If we did not re-sight families despite the extensive search of the study area, or we spotted the caring parent(s) alone without obvious signs of being alert and tending a brood, we considered the brood as failed.

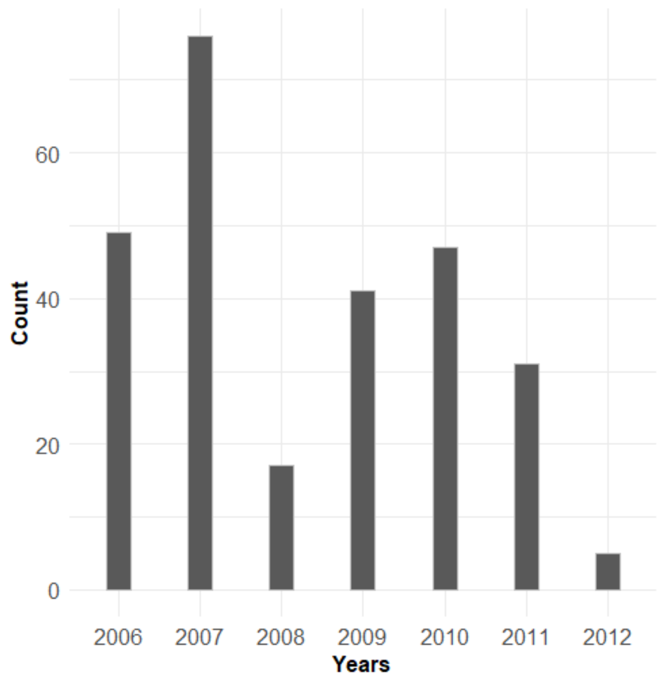


**Figure S1.** Number of Snowy Plover broods at Ceuta in the final data set during each study year from 2006 until 2012.

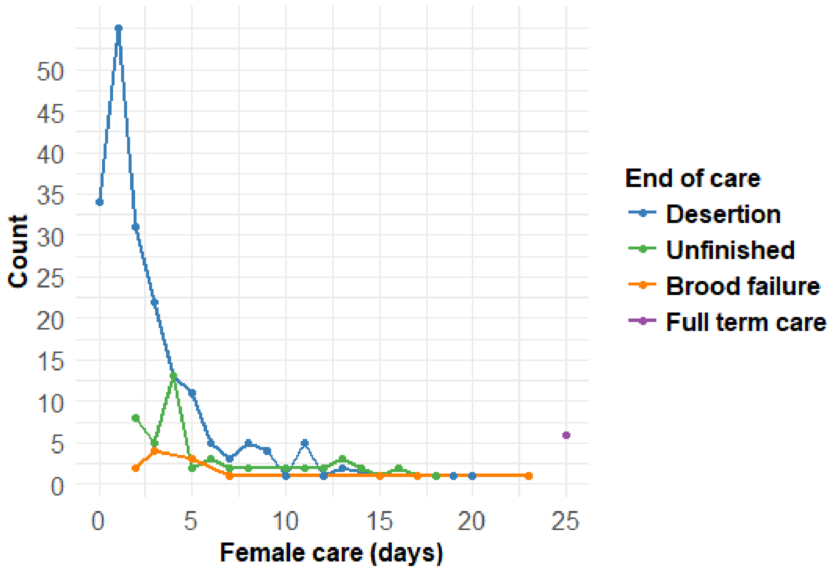


**Figure S2.** Length of female brood care in relation to brood fate in 268 Snowy Plover broods monitored at Ceuta.

I. **Female brood care**

**Reproductive success and breeding effort**

We calculated 'reproductive success' as the total number of fledglings for a certain female parental care strategy for a certain season. Many females likely disperse to different breeding sites between breeding attempts. Here, we compared the local 'breeding effort' of females that stayed for at least two breeding attempts or provided full term care in Ceuta. We defined the breeding effort as the number of days that we observed caring and deserting females during reproduction at Ceuta. The breeding effort represents the cumulated number of days of egg formation and incubation (3 + 25 days) and the days from hatching of the first chick to the end of care of the last brood. Therefore, it also includes the time that the female spent on finding a new mate after desertion (Figure S3). In the full term care group, all females had only one breeding attempt, whereas in the deserting group all females had two breeding attempts with different males. We compared breeding effort (in days) of the two groups using a Linear Model with log transformation of the response with *type of care* as predictor. We compared reproductive success (total number of fledglings) between the two groups using a Generalized Linear Model (glm) with Poisson error distribution.

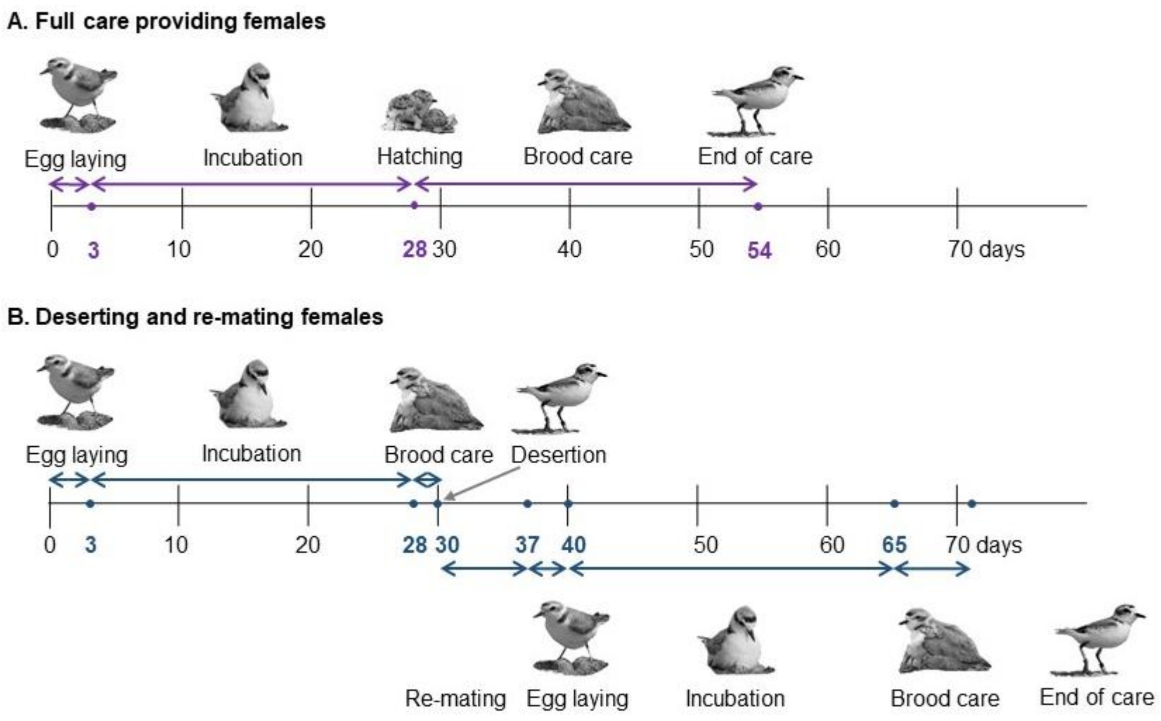


**Figure S3.** Examples for calculating breeding effort (length of care in days) in full term care providing (n = 6) (A) and deserting and re-mating (n = 9) females (B).

Deserting females that re-mated locally at our breeding site and successfully hatched chicks from both attempts (n = 9) did not have higher reproductive success than first time caring females (n = 6) (LM: Estimate: -0.09 ± 0.34, CI: -0.78 – 0.56, p = 0.788, Figure S4A). Both groups fledged on average 2.5 chicks. However, deserting females showed a higher variance in reproductive success than caring females with the most successful female having up to five fledglings.

The breeding effort of deserting females was substantially higher than the one of caring females (Figure 4B). Deserting females spent on average two and a half weeks (33%) longer at reproduction than females that cared for their chicks until fledging (GLM: Estimate: -0.28 ± 0.03, CI: -0.35 – -0.21, p < 0.0001).

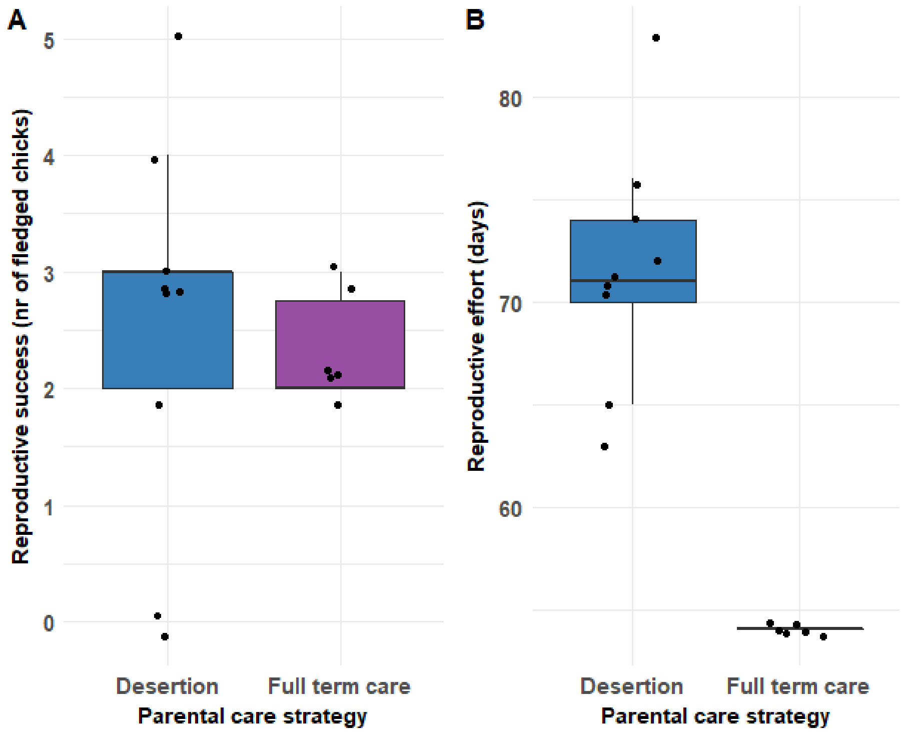


**Figure S4.** Reproductive success (A) and breeding effort (B) of deserting (n = 9) and full term caring (n = 6) Snowy Plover females at Ceuta.

### II. Predictors of the length of female brood care

*Detailed description of the predictors - Hatching date* provides a measure for the progress of the breeding season and we predicted that females would be more likely to stay and care for the chicks later in the season as re-mating opportunities diminish (Székely & Cuthill, 2000). We determined *hatching date* as relative hatching date, which is the z-transformation of the Julian hatching dates for each year. We calculated relative hatching date by using all broods with available hatching date information. As a measure of male size we used *male tarsus length*, the mean length of left and right tarsus of the male parent. *Male tarsus length* provides a measure of the size, thus protecting ability of the mate. We predicted that females would be more likely to care longer when paired to a smaller male that might not be as successful to protect the brood from conspecifics as a larger one. As another set of quality measures we calculated *male*, *female* and *chick condition* using the scaled mass index method by Peig and Green (Peig & Green, 2009). Positive values refer to individuals with better than the population average condition and negative values refer to individuals with worse than average condition. *Chick condition* was calculated using the hatching mass and size values of the chick in best condition in a brood, which highly correlated with the mean condition value of the brood. Similarly to *male tarsus length* we predicted that females would stay longer with males in lower condition. Females in lower condition should desert earlier to achieve high survival. We predicted that females would faster desert chicks with better condition as they have higher survival prospects than chicks with worse condition that might require more care. Alternatively, females might stay and care longer for chick in better condition as their survival prospects (and hence their reproductive value) are higher. *Present brood size* and *brood age* are two dynamic variables whose values were determined for every day over the brood care period. *Present brood size*, the brood size on a given day, allowed us to test whether females would take into account the present number of their chicks when deciding whether to desert or continue to care. We predicted that females would be more likely to continue to care for bigger than smaller broods. Finally, for *brood age* we predicted that older broods are more likely to be deserted as chicks are more independent (Currie et al., 2001; Gratto-Trevor, 1991). However, this relationship may be modified by seasonality. Because of reduced re-mating opportunities, some females especially at the end of the season may stay longer and care until fledging. Therefore, we also included the second polynomial of brood age (*brood age*$^2$).

**Effect of manipulation on the probability of care**

Brood manipulation experiments did not affect female *probability of care* (Figure S5a). Note that the effect of other predictor variables is nearly unchanged when comparing models with or without manipulation term (Figure 1a vs Figure S5a). However, removing all manipulated broods entirely from the analysis resulted in larger CrIs with the consequence that the effect of *brood age* on *probability of care* was statistically unclear using the reduced dataset (n = 155, Figure S5b). By contrast, *present brood size* and *hatching date* remained strong predictors of female *probability of care*.

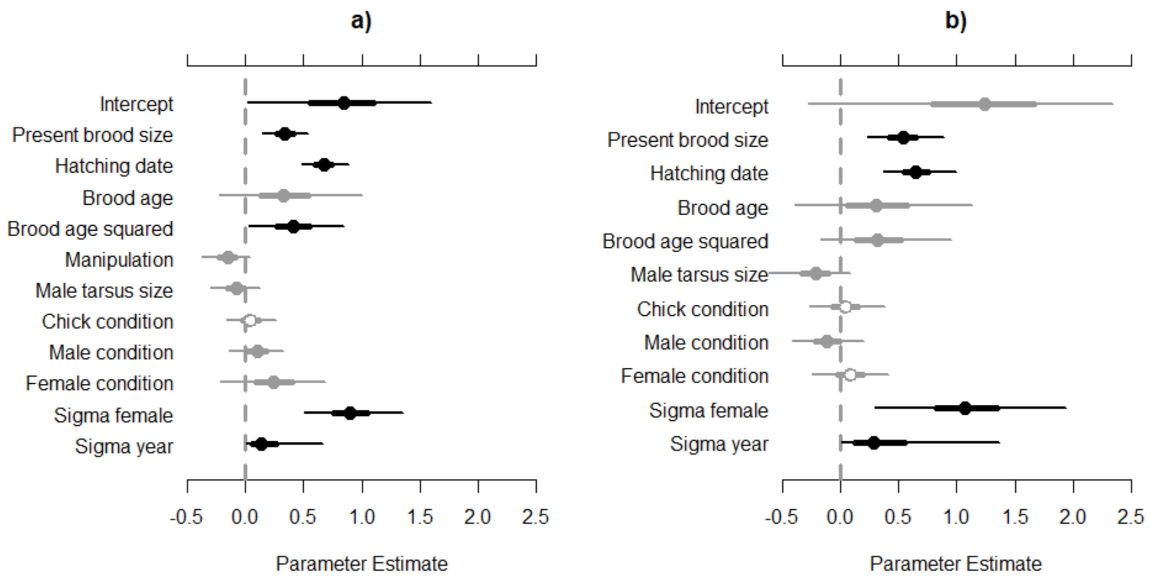


**Figure S5.** Effect plot of the predictors of female probability of care; a) controlling for experimental manipulations or b) removing all manipulated broods.

### III. Termination of care and chick mortality

Co-occurrence of desertion and chick mortality was influenced by the *present brood size* at termination of care. When a brood had one chick nearly all females deserted before that chick died or fledged (co-occurrence for broods with one chick: 7%, $p = 0.599$) meaning that there was little co-occurrence of chick death and termination of care. However, co-occurrence became statistically clear in broods with more than one chick at the time of termination of care (two chicks: 24%, $p < 0.001$; three chicks: 32%, no p value as model did not converge since *all* chick death events coincided with female desertion; >three chicks: 33%, no p value as model did not converge since *all* chick death events coincided with female desertion, Table S1, Figure S6).

| **Type of female care** | **Co-occurrence of chick mortality according to brood size** | | | | | | | |
|---|---|---|---|---|---|---|---|---|
| | 1 chick (n=56) | | 2 chicks (n=98) | | 3 chicks (n=56) | | 4-5 chicks (n=6) | |
| | No (0) | Yes (1) | No (0) | Yes (1) | No (0) | Yes (1) | No (0) | Yes (1) |
| Full term (0) | 1 | | 3 | | 2 | | 0 | |
| Terminated (1) | 51 | **4** | 71 | **24** | 36 | **18** | 4 | **2** |

**Table S1.** Co-occurrence of female care termination and chick death in 216 Snowy Plovers broods according to *present brood size* at care termination. ‘Full term’ refers to females that cared until their chicks fledged. ‘Terminated’ includes deserting females and those whose broods failed before the chicks fledged. Bold number indicates the number of broods for which chick mortality and care termination co-occurred on the same day.

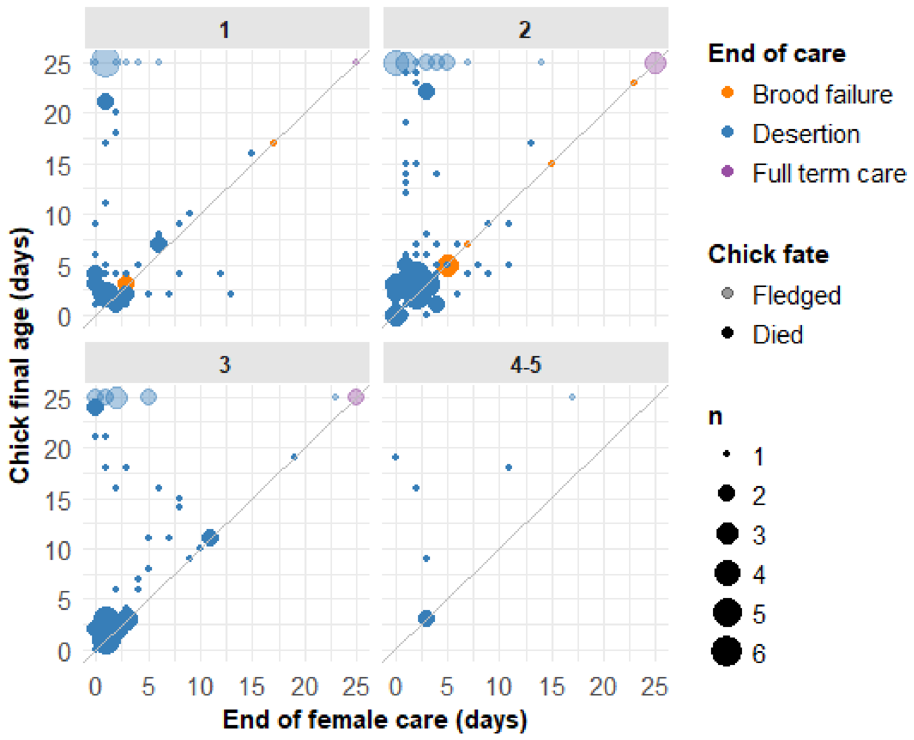


**Figure S6.** Chick survival in relation to length of female care for 182 Snowy Plover broods with known dates of chick death events according to *present brood size*. Only one chick per brood, whose fate was most closely associated with the end of female care is plotted. Dot sizes refer to number of broods. Coordinate (0,0) indicate the day of hatching and hence the start of female brood care. The diagonal line indicates co-occurrence of chick fate and female care termination on the same day. Below the diagonal are broods where the female continued to care after one chick had died, above the diagonal are broods where the female deserted before the first chick died.

**Chick survival in broods with or without co-occurrence of desertion and chick death**

We compared chick survival according to whether the chick came from a brood in which mortality and desertion co-occurred or not. We reasoned that broods in which chick mortality and desertion co-occur might indicate low survival prospects of these brood in general. Different survival between deserted broods with and without co-occurrence may also suggest that chick mortality is not simply a consequence of desertion of the female.

For the chick survival analysis we only included chicks from deserted broods in which at least one chick died to control for the statistical bias caused by the unavoidable chick mortality in the broods with co-occurrence. We applied mixed-effects cox models with the '*coxme*' function from the R package *coxme* and survival (T. M. Therneau, 2007; Terry M Therneau, Grambsch, & Pankratz, 2003). In the models we included the following predictors: *co-occurrence* ('Yes' or 'No'), *length of female care* (days of female presence) and individual *chick condition* (see calculation above). Furthermore, we added *Brood ID*, *Male ID* and *Year* as random variables (n = 345 chicks of 144 broods).

*Co-occurrence* and *length of female care* had a clear effect on chick survival, whilst individual chick condition did not influence survival (*co-occurrence*: coef = 0.73, p < 0.001; *length of female care:* coef = -0.09, p = 0.029, *chick condition*: coef = -0.02, p = 0.92; Figure S7).

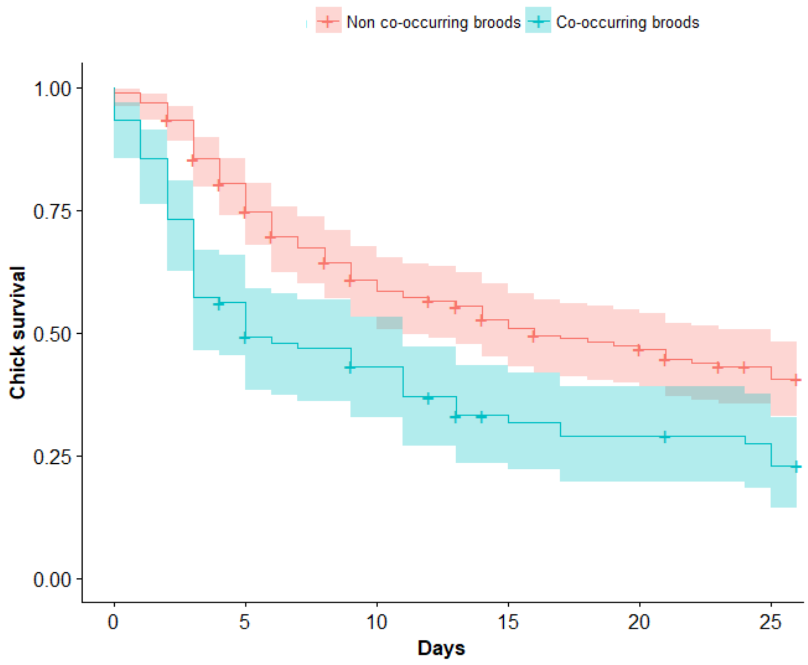


**Figure S7:** Kaplan-Meyer survival curves of Snowy Plover chicks from broods in which desertion and mortality co-occurred on the same day (n = 91) or not (n = 254).

**References:**

Cruz-López, M., Eberhart-Phillips, L. J., Fernández, G., Beamonte-Barrientos, R., Székely, T., Serrano-Meneses, M. A., & Küpper, C. (2017). The plight of a plover: Viability of an important snowy plover population with flexible brood care in Mexico. *Biological Conservation, 209*, 440-448. doi:https://doi.org/10.1016/j.biocon.2017.03.009

Currie, D., Valkama, J., Berg, Å., Boschert, M., Norrdahl, K., Häninnen, M., . . . Hemminki, O. (2001). Sex roles, parental effort and offspring desertion in the monogamous Eurasian Curlew Numenius arquata. *Ibis, 143*(3), 642-650. doi:10.1111/j.1474-919X.2001.tb04892.x

Dos Remedios, N., Székely, T., Küpper, C., Lee, P. L., & Kosztolányi, A. (2015). Ontogenic differences in sexual size dimorphism across four plover populations. *Ibis, 157*(3), 590-600.

Gratto-Trevor, C. (1991). Parental care in semipalmated sandpipers Calidris pusilla: brood desertion by females. *Ibis, 133*(4), 394-399.

Peig, J., & Green, A. J. (2009). New perspectives for estimating body condition from mass/length data: the scaled mass index as an alternative method. *OIKOS, 118*(12), 1883-1891.

Plaschke, S., Bulla, M., Cruz-Lopez, M., del Ángel, G., & Küpper, C. (2019). Nest initiation and flooding in response to season and semi-lunar spring tides in a ground-nesting shorebird. *Frontiers in Zoology, 16*(1), 11. doi:https://dx.doi.org/10.1186/s12983-019-0313-1

Székely, T., & Cuthill, I. C. (1999). Brood desertion in Kentish plover: the value of parental care. *Behavioral Ecology, 10*(2), 191-197.

Székely, T., & Cuthill, I. C. (2000). Trade-off between mating opportunities and parental care: brood desertion by female Kentish plovers. *Proceedings of the Royal Society of London Series B, 267*(1457), 2087-2092.

Therneau, T. M. (2007). On mixed-effect Cox models, sparse matrices, and modeling data from large pedigrees. *http://mayoresearch.mayo.edu/mayo/research/biostat/upload/kinship.pdf*. Retrieved from http://mayoresearch.mayo.edu/mayo/research/biostat/upload/kinship.pdf

Therneau, T. M., Grambsch, P. M., & Pankratz, V. S. (2003). Penalized survival models and frailty. *Journal of computational and graphical statistics, 12*(1), 156-175.
